# Control of neocortical memory by long-range inhibition in layer 1

**DOI:** 10.1101/2022.02.07.479360

**Authors:** Anna Schroeder, M. Belén Pardi, Joram Keijser, Tamas Dalmay, Erin M. Schuman, Henning Sprekeler, Johannes J. Letzkus

## Abstract

Mounting evidence identifies layer 1 (L1) as a central site of memory in sensory neocortex. While this work revealed plasticity in several excitatory brain-wide afferent systems, the existence, connectivity and memory-related signaling of long-range inhibitory input to L1 remains elusive. We report that inhibitory afferents from zona incerta project specifically to auditory cortex L1, where they connect selectively to interneurons to disinhibit the cortical circuit and facilitate behavioral memory. Chronic calcium imaging of these synapses identifies a balanced form of plasticity that develops rapidly during threat learning and is characterized by the *de novo* appearance of negative stimulus responses which transmit most information. Our results therefore pinpoint malleability of long-range (dis)inhibitory afferents to L1 as a key factor for the exquisite computational flexibility of this unique layer.

## Main Text

Sensory neocortex constantly constructs new memories. The underlying changes in information processing are driven by dynamic interactions between different neuron types, and depend on plasticity of both synaptic communication and network structure. However, localizing the plastic substrate of memory in dense and intricate neocortical circuits is often a major challenge. Recent evidence indicates that the outermost layer of neocortex (L1) is a major nexus for learning and memory (Shin et al., 2021). Different from all other layers, L1 is devoid of pyramidal neuron (PN) somata, and is instead comprised of PN dendrites and a unique set of interneurons (INs). These circuit elements receive long-range excitatory (LRE) afferents from a variety of memory-related areas (Doron et al., 2020; Gambino et al., 2014; Makino and Komiyama, 2015; Pardi et al., 2020), which are considered the dominant factor for the highly experience-dependent signaling within L1 itself (Letzkus et al., 2015; Roelfsema and Holtmaat, 2018; Shin et al., 2021). However, in parallel to these intensely investigated systems, cortical structures contain a sparser and much less understood complement of long-range inhibitory (LRI) projections (Kaifosh et al., 2013; Melzer and Monyer, 2020). Therefore, our objectives were to determine whether L1 of sensory neocortex receives LRI, and to identify its potentially unique contributions to memory.

The subthalamic zona incerta (ZI) is an enigmatic inhibitory brain region that integrates multisensory information and regulates a wide range of behaviors including learning (Ahmadlou et al., 2021; Li et al., 2021; Liu et al., 2017; Masri et al., 2009; Zhang and van den Pol, 2017; Zhou et al., 2018). Since this area orchestrates cortical development in neonatal mice through LRI afferents to L1 (Chen and Kriegstein, 2015), we asked whether a similar projection is present in adult auditory cortex (ACx). Indeed, anterograde tracing from GABAergic neurons after injection of a conditional adeno-associated viral vector (AAV) into the ZI of GAD2-IRES-Cre mice (Fig. 1, A and B) revealed a robust projection. These afferents are strongly enriched in L1 of area AuV (Fig. 1, C and D), the most temporal region of ACx which is also critical for auditory threat memory (Dalmay et al., 2019). To determine the proportion of ACx-projecting ZI neurons that are inhibitory, we injected retrograde tracers in GAD2-nuclear-mCherry mice (Fig. 1E). This revealed that a subset of overwhelmingly GABAergic neurons located in the ventral ZI projects to ACx (Fig. 1, F and G). Is this novel LRI projection a general feature of sensory areas of neocortex? Triple retrograde tracing from visual cortex (VCx), somatosensory cortex (SSCx) and ACx (Fig. 1H) indicates that the vast majority of labeled neurons project to ACx (Fig. 1I), suggesting that this pathway may preferentially contribute to auditory behavior. Together, these data identify the ZI as a major source of LRI in L1 of the ACx.

**Fig. 1.**
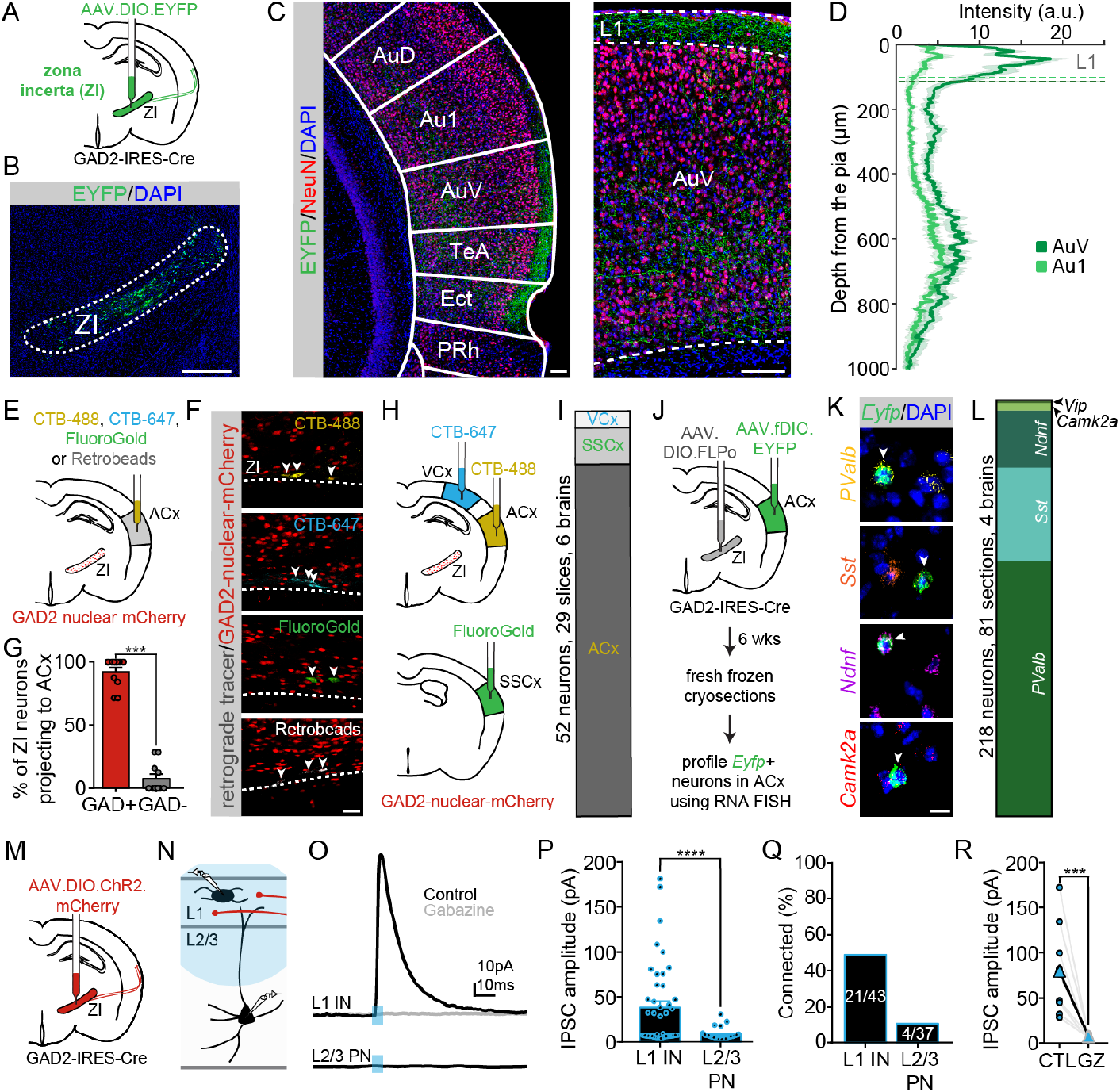
Long-range inhibition from zona incerta preferentially targets interneurons in auditory cortex. (**A**) Schematic for anterograde tracing. (**B**) Viral expression of EYFP in GABAergic ZI neurons. (**C**) Left: ZI axons display a mediotemporal density gradient and preferentially target L1. Right: Close-up of AuV. (**D**) Axon density across cortical depth in primary and secondary ACx (Au1 & AuV, n = 12 sections, 5 mice). L1 width; dotted lines. (**E**) Schematic for retrograde tracing from ACx. (**F**) Example images of ACx-projecting GABAergic ZI neurons (arrowheads). (**G**) The large majority of ACx-projecting ZI neurons is GABAergic (42 sections, 11 mice). (**H**) Schematic for triple retrograde tracing from ACx, SSCx and VCx. (**I**) Fraction of ZI neurons projecting to ACx, SSCx or VCx out of all cortex-projecting neurons (n = 52 neurons, 29 slices, 6 mice). ZI projects most strongly to ACx (**J**) Schematic for AAV1-mediated anterograde transsynaptic tracing combined with FISH for identifying postsynaptic targets. (**K**) Examples of *Eyfp*-positive (green) neurons expressing either *PValb* (yellow), *Sst* (orange), *Ndnf* (violet) or *Camk2a* (red). (**L**) ZI overwhelmingly targets interneurons expressing the markers *PValb, Sst* or *Ndnf* in ACx (n = 218 neurons, 81 sections, 4 mice). (**M**) Schematic for ChR2 expression in GABAergic ZI axons. (**N**) ZI axons were stimulated in acute brain slices by light pulses (5 ms) directed at the superficial layers of AuV while L1 INs or L2/3 PNs were recorded. (**O**) Example mean IPSC traces (blue: light pulse). (**P**) IPSC amplitudes for all recorded neurons (both connected and non-connected). ZI inputs to L1 INs (38.6 ± 6.9 pA n = 43 L1 INs, 18 mice) are stronger than for L2/3 PNs (7.7 ± 0.9 pA, n = 37 L2/3 PNs, 18 mice). (**Q**) Percentage of connected neurons. (**R**) Application of Gabazine (GZ) to connected L1 INs (CTL) abolishes light-evoked responses (n = 10 L1 INs). Data represent mean ± SEM. *** p<0.001, **** p<0.0001. Scale bars: (B) 500 μm, (C) 100 μm, (F) 40 μm, (K) 10 μm. Statistics: (G) Wilcoxon matched-pairs signed rank test; (P) Mann Whitney test; (R) 2-tailed paired t-test.

How these inputs control the local circuit depends on their targets. To determine the identity of the postsynaptic neurons, we employed AAV1-mediated anterograde transsynaptic tracing (Zingg et al., 2020) in GAD2-IRES-Cre mice in combination with fluorescent *in situ* hybridization (Fig. 1J, see Methods). Strikingly, >97% of profiled neurons were inhibitory INs expressing the markers *Pvalb, Sst* or *Ndnf*, whereas only few excitatory *Camk2a* or disinhibitory *Vip* neurons were found (Fig. 1, K and L). Functional validation of the observed connectivity was carried out using channelrhodopsin-2 (ChR2)-assisted circuit mapping (Petreanu et al., 2009) in acute brain slices while recording from ACx L1 INs and L2/3 PNs (Fig. 1, M and N). Light stimulation of superficial layers caused inhibitory currents in L1 INs with approximately fivefold higher amplitudes (Fig. 1, O and P) and with greater probability (Fig. 1Q) than in L2/3 PNs. These inputs are mediated by GABAA receptors (Fig. 1R), and display longer latencies in L2/3 PNs than in L1 INs, consistent with innervation of distal PN dendrites in L1. Collectively, these results indicate that LRI ZI afferents predominantly inhibit INs in ACx, whereas direct inhibition of PN dendrites in L1 is considerably weaker. Importantly, the resulting net disinhibition of the local circuit has been identified as a conserved processing motif enabling network plasticity during learning and memory (Basu et al., 2016; Letzkus et al., 2015).

On top of its output connectivity, a major determinant of the *in vivo* function of the ZI-ACx pathway is the inputs it receives. To specifically identify the brain-wide inputs to ZI neurons with a direct projection to the ACx, we used a combination of retrograde and transsynaptic rabies tracing (Fig. 2, A and B). This uncovered a number of input sources spanning across the neuroaxis (Fig. 2, C and D), including the midbrain, striatum, thalamus and cortex. These data thus reveal that ZI sends integrated information from diverse upstream areas to ACx. Of note, several of these input regions such as the higher-order auditory thalamus, central amygdala and periaqueductal gray have been implicated in threat learning (Herry and Johansen, 2014), suggesting that ACx-projecting ZI neurons may play a central role for this form of memory.

**Fig. 2.**
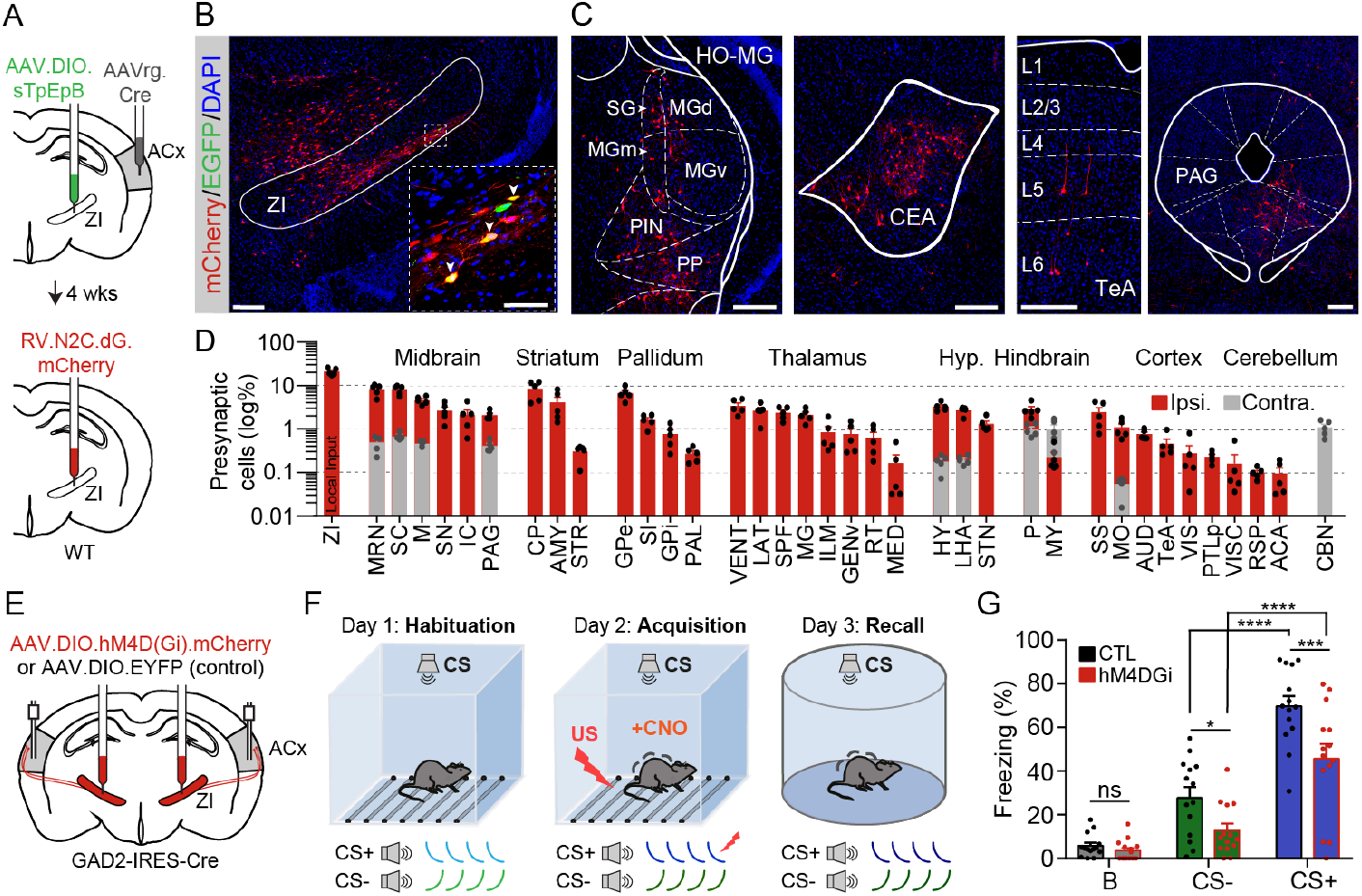
Zona incerta provides integrated information to auditory cortex that is essential for threat memory. (**A**) Schematic for identifying brain-wide monosynaptic inputs to ZI neurons that project to ACx. (**B**) Starter cells (EGFP+/mCherry+, arrowheads), presynaptic cells (mCherry+) and non-starter cells (EGFP+) in the ZI. (**C**) Representative presynaptic neurons (HO-MG, higher-order auditory thalamus; CEA, central amygdala; TeA, temporal association cortex; PAG, periaqueductal grey) in regions essential for auditory threat learning. (**D**) Distribution of presynaptic cells ipsilateral (red) and contralateral (grey) to injection (n = 5 mice, logarithmic scale, bars overlap). (**E**) Schematic for bilateral chemogenetic inhibition of ZI afferents in ACx by local agonist application via cannulae. (**F**) Discriminative auditory threat conditioning paradigm using complex CSs. CNO was infused prior to acquisition. (**G**) Freezing behavior during recall indicates reduced memory strength compared to controls for both the CS+ and the CS- (n = 14 mice per group). Abbreviations used in (C) and (D) are defined in table S1. Data represent mean ± SEM. n.s. p>0.05, * p<0.05, *** p<0.001, **** p<0.0001. Scale bars: (B) 200μm (inset, 50μm), (C, all bars) 200μm. Statistics: (G) Two-way RM ANOVA with Tukey’s multiple comparisons test between trial types and Sidak’s multiple comparisons test between animal groups.

To directly address whether inhibitory ZI afferents in ACx impact learning, we employed a form of discriminative threat conditioning (DTC) that critically depends on ACx function due to the use of complex conditioned stimuli (CSs, Fig. 2, E and F) (Ceballo et al., 2019; Dalmay et al., 2019). We expressed the chemogenetic inhibitor hM4DGi in GAD2-positive ZI neurons, and implanted bilateral cannulae over ACx (Fig. 2E). This permitted temporally and spatially controlled infusion of the ligand clozapine-N-oxide (CNO) in ACx prior to the memory acquisition session, during which one of two initially neutral CSs (CS+) was repeatedly paired with an unconditioned stimulus (US, a mild foot-shock), while the CS- was left unpaired. Silencing of LRI ZI axons in ACx during learning resulted in a robust memory deficit during recall, as evidenced by less freezing of the mice to both CSs (Fig. 2G). Conversely, there was no effect on acute freezing behavior during acquisition, on CS discrimination during recall, or on recall of contextual threat memory. These results therefore indicate that LRI ZI projections to ACx selectively control the formation of long-term auditory threat memory.

How may information transmitted by ZI afferents to ACx L1 contribute to associative memory? To address this, we performed chronic 2-photon imaging of ZI synaptic boutons expressing an axon-targeted calcium indicator (Broussard et al., 2018) in L1 of awake, head-fixed GAD2-IRES-Cre mice after identification of AuV using intrinsic imaging (Fig. 3A) (Abs et al., 2018; Dalmay et al., 2019; Pardi et al., 2020). This was combined with a novel DTC paradigm in which all phases occurred in head-fixation, enabling us to longitudinally track the responses of individual boutons prior to (habituation), during (acquisition) and after learning (recall, Fig. 3B to D) while using changes in eye pupil dilation in response to the CSs as an online readout of threat memory (Fig. 3D) (Abs et al., 2018; Dalmay et al., 2019; Pardi et al., 2020). Note that only boutons which could be identified in all three sessions were analyzed (Fig. 3C). Presentation of CSs in the habituation session elicited clear positive or negative responses in some boutons (Fig. 3, E to G), indicating that this non-canonical pathway transmits auditory information. Training a linear decoder to predict stimulus identity from the bouton response patterns furthermore showed that LRI ZI afferents are able to discriminate auditory stimuli above chance level at both the single-bouton (Fig. 3H) and population level (Fig. 3I). Given that the CSs largely overlap in frequency content, this suggests remarkably precise encoding of auditory information by this pathway. In addition, ZI boutons respond robustly to the mild tail-shock used as the US (Dolensek et al., 2020). These results uncover that LRI ZI afferents transmit information about both the auditory CSs and the aversive US to ACx.

**Fig. 3.**
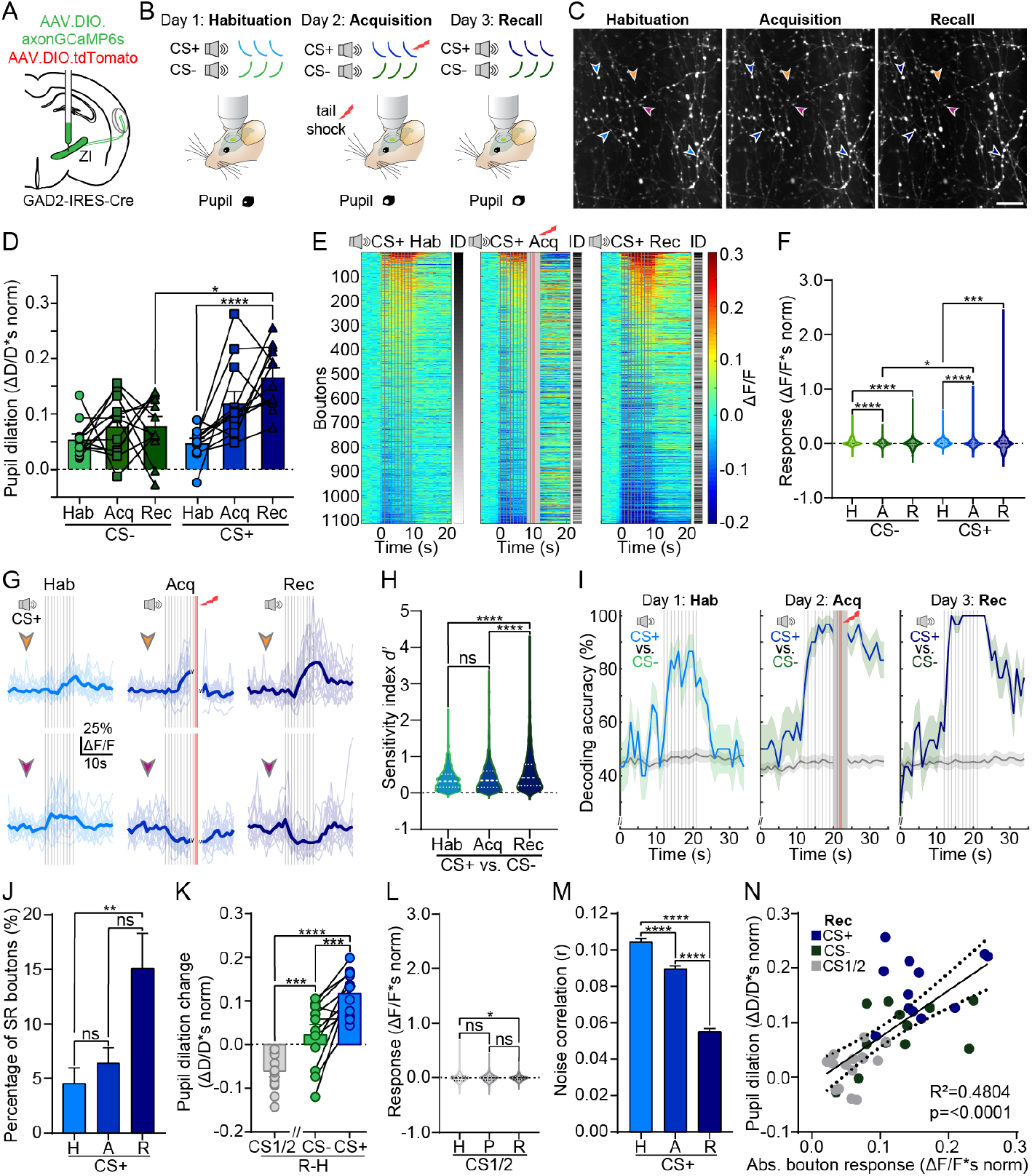
Robust response plasticity of zona incerta synapses in layer 1 upon learning. (**A**) Schematic for chronic *in vivo* 2-photon calcium imaging of GABAergic ZI boutons in ACx L1 through a cranial window. (**B**) Discriminative auditory threat conditioning in head-fixation with eye pupil dilation as an online memory readout. (**C**) Representative boutons across sessions. (**D**) Pupil dilation in response to the CSs across behavioral sessions (n = 12 mice). (**E**) Trial-averaged single bouton responses to the CS+ sorted by amplitude (gray scale: rank order of bouton ID, vertical lines: FM-sweeps (gray) and tail-shocks (red), light gray: blanked period during acquisition). (**F**) Response integral distribution broadens across the paradigm for both CSs (n = 1114 boutons, 12 mice). (**G**) Representative single trial (thin lines) and average (thick lines) CS+ responses illustrating excitatory and inhibitory potentiation of the boutons marked in (C). (**H**) Stimulus discriminability quantified for single boutons as the sensitivity index d’ (n = 1114 boutons). (**I**) Accuracy of bouton population decoding between CS+ and CS- increases across the paradigm (n = 1114 boutons, grey: chance level estimated by permuting stimulus labels). (**J**) Percentage of CS+ sound responsive (SR) boutons (n = 12 mice). (**K**) Pupil dilation change between habituation and recall in threat conditioned (n = 12 mice) and pseudoconditioned animals (n = 8 mice, CS1 and CS2 combined). (**L**) Same as in F for pseudoconditioned animals (n = 1070 CS1 and CS2 responses, 535 boutons, 8 mice). (**M**) Noise correlations of CS+ responses between all pairs of boutons (n = 76844 pair-wise correlations, 1114 boutons, 12 mice). (**N**) Correlation between pupil dilation and mean absolute bouton response in recall (n = 12 threat conditioned mice, 8 pseudoconditioned mice). Data represent mean ± SEM. n.s. p>0.05, * p<0.05, ** p<0.01, *** p<0.001, **** p<0.0001. H, Hab: habituation; A, Acq: acquisition; R, Rec: recall. Scale bar: (C) 20μm. Statistics: (D) RM One-way ANOVA with Sidak’s multiple comparisons test, (F, H, K, L, M) Friedman test with Dunn’s multiple comparisons test, (J) Ordinary one-way ANOVA with Sidak’s multiple comparisons test, (N) Pearson correlation, linear fit and 95% confidence bands.

On top of these sensory signals, ZI boutons also displayed pronounced plasticity of CS response patterns due to DTC. These changes were bidirectional, with some boutons developing strong positive responses (excitatory potentiation) while others started to display negative responses (inhibitory potentiation). Plasticity emerged during memory acquisition, and was even more robust in memory recall (Fig. 3, E to G). It materialized in all possible rearrangements, with either positive or negative responding boutons undergoing either excitatory or inhibitory potentiation. Moreover, the percentage of responsive boutons grew with learning (Fig. 3J), driven by the emergence of strong negative responses that were largely absent prior to learning. These plastic changes occurred for the CS+, and to a lesser extent for the CS-, consistent with the strength of threat memory to these stimuli (Fig. 3, D and K). In contrast, control animals that underwent non-associative pseudoconditioning (PC) and thus did not form a threat memory showed no bidirectional changes (Fig. 3, K and L), demonstrating that the plasticity seen in conditioned mice is specific for associative learning.

One notable consequence of the bidirectional changes is an increase in the inter-bouton standard deviation of responses, raising the question how this affects transmission of sensory information. Single bouton and population decoding analysis showed that stimulus discrimination by LRI ZI boutons improves with threat learning (Fig. 3, H and I), in line with the enhanced behavioral discrimination of CS+ and CS-, whereas both effects were absent in pseudoconditioned animals. We next analyzed trial-by-trial activity correlations between boutons, which can limit the content of information transfer about sensory stimuli. Threat memory acquisition resulted in a decrease in pairwise noise correlations between boutons which was not observed after PC, suggesting a learning-related increase in the amount of information transferred by this pathway (Fig. 3M). To elucidate how learning changes CS encoding in LRI ZI boutons at the population level, we computed the population vectors of responses in habituation and recall, and quantified the angle between these vectors. This revealed that DTC causes a robust change in the representation of both CS+ and CS-, at almost twice the angle observed in control mice after PC. Importantly, these changes caused the boutons to encode memory strength: The absolute response to a given CS in the recall session correlates with pupil dilation as an online readout of memory, while this relationship is absent in habituation (Fig. 3N). In conclusion, threat memory manifests as a balanced form of population plasticity in ZI afferents that improves CS discrimination and information transfer, and encodes the strength of the memory trace, in line with the observed essential function of this pathway for learning (Fig. 2G).

Comparable experiments on excitatory afferents to L1 have linked memory exclusively to potentiation of positive responses, i.e. a strengthening of synaptic transmission (Doron et al., 2020; Gambino et al., 2014; Makino and Komiyama, 2015; Pardi et al., 2020), indicating that the observed balanced and bidirectional plasticity may be a novel hallmark of LRI. To address the functional consequences of the *de novo* appearance of negative stimulus responses, we segregated the boutons into two groups based on whether they showed positive (PR; average ΔF/F*s > 0) or negative responses (NR; average ΔF/F*s < 0) to the CS in recall (Fig. 4, A and B). This revealed that both groups display positive mean responses during habituation, and that the change during memory acquisition is predictive of their responses during recall (Fig. 4, C and D). Boutons in either group also responded to the US with both positive and negative transients, but no relationship between these post-shock responses and the CS response during acquisition or recall was identified. To address the dynamics of learning-related plasticity, we plotted the responses across trial bins (Fig. 4, E and F). This uncovered a striking dichotomy between PR and NR boutons during the acquisition session: While the main effect of learning for PR boutons is a stabilization of responses across trials, NR synapses display a rapid switch from initially positive to negative responses over just a few presentations of the CS+/US compound. This effect parallels the evolution of behavioral online learning, and highlights the central role of negative responses for acute encoding of memory by LRI afferents. Conversely, both PR and NR bouton responses were robustly potentiated in the memory consolidation time window between acquisition and recall, enabling both populations to encode memory strength in the recall session. These results support the idea that LRI ZI boutons engage in two plasticity regimes with distinct temporal dynamics and response properties, which unfold during memory acquisition, persist in recall (Fig. 4, A to F), and are absent in control animals.

**Fig. 4.**
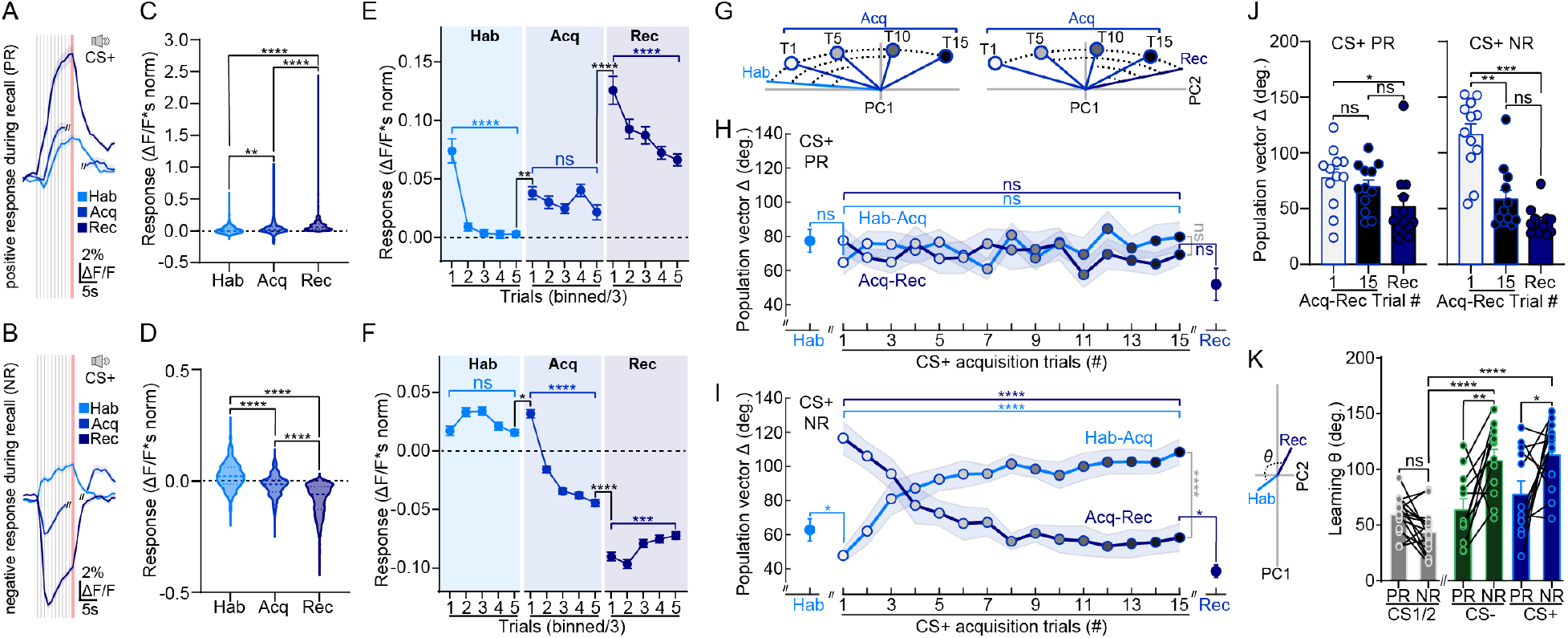
Rapid inhibitory potentiation drives learning. Boutons were segregated based on whether they displayed a positive response (PR, n = 531 boutons) or a negative response (NR, n = 583 boutons) during threat memory recall (n = 12 mice). (**A**) Mean CS+ responses of PR boutons across the paradigm. (**B**) Same as A for NR boutons. (**C, D**) Integral of CS+ responses (PR, n = 531 boutons; NR, n = 583 boutons). (**E**) CS+ response integral over trials (every three binned) for PR boutons across the paradigm (n = 531 boutons). (**F**) Same as E for NR boutons (n = 583 boutons). Note rapid appearance of negative responses during acquisition. (**G**) Schematic for quantification of learning dynamics as the angle between CS+ population response vectors in each acquisition trial relative to habituation (left, light blue) or recall (right, dark blue). Values for habituation and recall represent trial-to-trial variability within those sessions. (**H**) PR boutons show no dynamic change across learning (n = 12 mice). (**I**) NR boutons display rapid learning dynamics (n = 12 mice). (**J**) Angle between CS+ population response on first and last acquisition trials relative to recall (PR, left; NR, right) indicates greater changes for NR boutons (n = 12 mice). (**K**) Angle between average habituation and recall population vectors for PR and NR boutons indicates greater contribution of negative responses to threat memory (n = 12 threat conditioned mice; 16 CS1 and CS2 representations from 8 pseudoconditioned mice). Data represent mean ± SEM. n.s. p>0.05, * p<0.05, ** p<0.01, *** p<0.001, **** p<0.0001. Vertical lines in A, B: FM-sweeps (gray), tail-shocks (red). Statistics: (C, D) Friedman test with Dunn’s multiple comparisons test, (E, F) Two-way ANOVA, with Sidak’s multiple comparisons test, (H, I) Two-way ANOVA with Sidak’s multiple comparisons test or 2-tailed paired t-test, (J) RM One-way ANOVA with Tukey’s multiple comparisons test, (K) Ordinary one-way ANOVA, with Sidak’s multiple comparisons test.

To further dissect the potentially unique function of LRI-specific NR boutons, we addressed how they contribute to learning-related plasticity of CS encoding at the population level by computing the angle between individual acquisition trial population vectors and the average population vector during either habituation or recall (Fig. 4G). Over consecutive CS+/US pairings, the angle between habituation and acquisition responses successively increased for NR boutons, while it simultaneously decreased between acquisition and recall (Fig. 4, H to J). These changes occur rapidly: within only five acquisition trials, the CS representation has become more similar to the final pattern that will emerge in recall than to the original one in habituation. In stark contrast, PR boutons showed no change in CS encoding during learning (Fig. 4, H and J). These results are largely mirrored, albeit with lower magnitude, for the CS-, and do not occur in control mice. Do either PR or NR boutons make a privileged contribution also to long-term memory during recall? Reanalysis of the population level changes between habituation and recall, this time for PR and NR boutons separately, reveals a greater degree of plasticity in NR boutons for both CS+ and CS- relative to the PR group (Fig. 4K). In conclusion, rapid inhibitory potentiation is the major driver converting population response patterns from encoding neutral sensory information during habituation to representation of stimuli with learned relevance during recall.

Despite the therapeutic capacity of the ZI (Ossowska, 2020), and its central involvement in numerous behaviors including sleep (Liu et al., 2017), feeding (Zhang and van den Pol, 2017), pain (Masri et al., 2009), novelty-seeking (Ahmadlou et al., 2021), anxiety (Li et al., 2021) and threat memory (Zhou et al., 2018), the function and information content of its projection to neocortex has remained elusive. This study identifies LRI ZI afferents as a source of remarkably precise auditory information and reinforcement signals in ACx which selectively connect to INs, widely integrate information from across the neuroaxis, are essential for associative memory and undergo complex, bidirectional plasticity during and after learning. Importantly, the rapid appearance of negative stimulus responses in approximately half of the ZI boutons after just a few conditioning trials contrasts markedly with results from several different LRE afferents in L1, for which exclusively positive transients and excitatory potentiation have been observed during memory acquisition and expression (Doron et al., 2020; Gambino et al., 2014; Makino and Komiyama, 2015; Pardi et al., 2020; Shin et al., 2021), pinpointing a potentially unique attribute of LRI in L1. These bidirectional changes encode the strength of the memory trace and improve stimulus discrimination in the absence of large effects on mean responses, uncovering that on top of their well investigated function as a disinhibitory gate for plasticity induction that has been described in the hippocampal formation (Basu et al., 2016), information transmitted by LRI projections is itself subject to computationally rich modifications during learning. The bidirectional implementation of this plasticity may serve to improve both the dynamic range and the metabolic efficiency of information transfer to cortical circuits. Given how little we know yet, the biological implications of these mechanisms and their involvement in other forms of learning and behavior need to be further investigated.

LRI ZI projections likely contribute to associative learning via two distinct operations: First, acute disinhibition recruited by the US can instruct plasticity induction in the local circuit (Letzkus et al., 2015). Second, changes in CS encoding by ZI afferents will directly contribute to the modified representation of these stimuli in ACx (Ceballo et al., 2019; Dalmay et al., 2019). An attractive possibility for the mechanistic implementation of these effects is that the ZI – via its profuse projections to numerous brain areas in addition to ACx (e.g. Trageser and Keller, 2004; Urbain and Deschenes, 2007) – serves to temporally coordinate and organize the activity patterns within this brain-wide network in a manner that enables memory formation and recall (Dejean et al., 2016). Notwithstanding the precise mechanistic realization, our multidisciplinary dissection of ZI afferents has begun to reveal the importance and unique attributes of neocortical LRI, and may also serve to motivate work on the contribution of further LRI systems (e.g. Saunders et al., 2015) to the computational power and flexibility of neocortical circuits.

## Acknowledgments

We thank members of the Letzkus, Sprekeler and Schuman labs, M.S. Fustinana Gueler and P. Tovote for discussions; N. Dolensek, S. Junek, F. Vollrath, F. Kretschmer, and G. Tushev for technical assistance; G. Keller for technical comments; and L. L. Looger, J. Akerboom, D. S. Kim, the GENIE Project at Janelia Farm, K. Deisseroth, E. S. Boyden, H. Zeng, L. Tian, L. Zhang, J. M. Wilson, B. Roth and K.-K. Conzelmann for generously sharing reagents.

## Funding

JJL: Max Planck Society, German Research Foundation (LE 3804/3-1, LE 3804/4-1, LE 3804/7-1). AS: Marie Skłodowska-Curie Fellowship (840701), EMBO Long-Term Fellowship (ALTF 882-2018), Alexander von Humboldt Fellowship.

## Author contributions

Conceptualization: AS, JJL. Project administration: AS, JJL. Writing: AS, JJL with input from all authors. Investigation: AS, TD. Formal analysis: AS, JK, HS. Methodology: AS, JK, MBP, HS, EMS, JJL. Software: JK, HS. Visualization: AS. Supervision: JJL.

## Competing interests

Authors declare no competing interests.

## Code availability

Code for analysis and computational work is available on GitHub or GitLab (Castaño-Díez, 2018; Keijser, 2022; Kretschmer, 2020a, b; Kretschmer and Castaño-Díez, 2020; Kretschmer and Unzué, 2020).

## Materials and Methods

### Animal model

All animal procedures were executed in accordance with institutional guidelines and approved by the prescribed authorities (*Regierungspräsidium* Darmstadt, approval numbers F126/1027 and F126/2000). Adult (>P35) homozygous GAD2-IRES-Cre mice (Gad2^tm2(cre)Zjh/J^, JAX stock #010802, The Jackson Laboratory) (Taniguchi et al., 2011) and GAD2-T2a-NLS-mCherry mice (Gad2^tm1.1Ksvo^, JAX stock #023140, The Jackson Laboratory) (Peron et al., 2015) were used. Animals were housed under a 12 h light/dark cycle, and provided with food and water *ad libitum*, except for water restriction periods in mice trained for head-fixation (body weight loss <15%). After surgical procedures, mice were individually housed. In behavioral experiments, only male mice were used. Research was conducted following the ARRIVE guidelines.

### Surgery

In all cases, mice were anesthetized with isoflurane (induction: 4%, maintenance: 2%) in oxygen-enriched air (Oxymat 3) and fixed in a stereotaxic frame (Kopf Instruments). Body temperature was maintained at 37.5 °C by a feedback controlled heating pad (FHC). Analgesia was provided by local injection of Ropivacain under the scalp (Naropin) and i.p. injection of metamizol (200 mg/kg, Novalgin, Sanofi) and meloxicam (2 mg/kg, Metacam, Boehringer-Ingelheim). For surgeries involving chronic window implantation, buprenorphine (i.p injection, 0.1 mg/kg, cp-pharma) was used instead of metamizol. Adeno-associated viral vectors (AAVs) or retrograde tracers were injected from glass pipettes (P0549, Sigma) connected to a pressure ejection system (PDES-02DE-LA-2, NPI). For ZI, 200 nl of AAV was injected at −1.94 mm posterior and ±1.7 mm lateral from bregma, and −4.15 mm from the cortical surface. For AAV injections in ACx, 500 nl of AAV was injected at −2.54 mm posterior and +4.6 mm lateral from bregma, and −1 mm from cortical surface. For retrograde tracing experiments, 250nl of CTB-488 (#C34775, Thermo Fisher Scientific), CTB-647 (#C34778, Thermo Fisher Scientific) or 4% FluoroGold (Fluorochrome) were injected in ACx (same coordinates as for AAV), SSCx (−0.23 mm posterior, +4.0 mm lateral from bregma, and −0.75 mm from cortical surface) or VCx (−3.79 mm posterior, +3.0 mm lateral from bregma, and −0.3 mm from cortical surface) of GAD2-nuclear-mCherry mice. For AAV experiments, injection was followed by 5 weeks of expression, while for retrograde tracing, there was 1 week between injection and tissue collection. In surgeries for anterograde tracing experiments, GAD2-IRES-Cre mice were injected with AAV2/5-EF1a-DIO-EYFP-WPRE-hGH (Penn) in the ZI. For transsynaptic tagging experiments, GAD2-IRES-Cre mice were injected with AAV2/1-pEF1a-DIO-FLPo-WPRE-hGHpA (Addgene #87306-AAV1) in ZI, and in ipsilateral ACx with AAV2/5-EF1a-fDIO-EYFP-WPRE (UNC). For acute slice experiments, GAD2-IRES-Cre mice were injected with AAV2.5-Ef1a-dflox-hChR2(H134R)-mCherry.WPRE.hGH (Penn) in ZI. For rabies tracing, WT mice were first injected in the ZI with AAV-synP-DIO-sTpEpB (UNC), and in ACx with pENN-AAV-hSyn-Cre-WPRE-hGH (Addgene #105553-AAVrg) in an AAVrg backbone, and 6 weeks after expression, in ZI with RV-N2C-dG-mCherry-EnvA (kindly provided by Karl-Klaus Conzelmann). For chemogenetics experiments, GAD2-IRES-Cre mice were injected bilaterally with AAV2/5-hSyn-DIO-hM4D(Gi)-mCherry (UNC) in ZI. For the corresponding control group, either C57BL/6J or GAD2-IRES-Cre mice were injected bilaterally with AAV2/5-CamKII0.4-eGFP-WPRE-rBG (Penn) or AAV2/5-EF1a-DIO-EYFP-WPRE-hGH (Penn), respectively, in ZI. For both experimental and control animals, bilateral cannulae (Bilaney, #C315G/Spc, #C315I/Spc and #C315DC/Spc) were then implanted over ACx (−0.7mm from cortical surface). In surgeries for calcium imaging experiments, GAD2-IRES-Cre mice were injected in the ZI with a 5:1 mix of AAV2/5-hSynapsin1-Flex-axon-GCaMP6s (Addgene, #112010-AAV5) and AAV2/9-FLEX-tdTomato (Addgene, #28306-AAV9). For chronic window implantation, a craniotomy was performed over ACx with a biopsy punch (Integra Miltex), and covered by a custom made window (a round cover glass glued with Norland optical adhesive #81 to a section of hypodermic tubing of outer diameter 3 mm). The window and a custom-made titanium head plate were fixed using Cyanoacrylate glue (Ultra Gel, Henkel) and dental cement (Paladur, Heraeus). The glass window was protected with silicone (Kwik-Cast). Calcium imaging was performed >5 weeks after surgery.

### Transsynaptic tagging with fluorescent in situ hybridization (FISH)

GAD2-IRES-Cre mice were injected with AAV2/1-pEF1a-DIO-FLPo-WPRE-hGHpA (Addgene #87306-AAV1) in ZI, and in ipsilateral ACx with AAV2/5-EF1a-fDIO-EYFP-WPRE (UNC). 6 weeks post-injection, mice were anesthetized with isofluorane, sacrificed and the brains were then dissected, embedded in Tissue-Tek O.C.T. compound (Sakura) and frozen in isopentane at −55 to −60°C. 16 μm-thick sections from these fresh frozen brains were prepared using a cryostat (Leica) and mounted on SuperFrost Plus microscope slides (Thermo Scientific). Sections were screened for fluorescence in auditory cortex (Zeiss Axio Zoom) and then stored at −80°C until FISH was performed using the RNAscope Fluorescent Multiplex Reagent Kit (#320850, Advanced Cell Diagnostics). Heating steps were performed using the HybEZTM oven (Advanced Cell Diagnostics). Tissue sections were treated with pretreatment solutions and then incubated with RNAscope probes (EGFP (which also labels EYFP), #400281; EGFP-sense-C2, #409971-C2; Mm-Camk2a-C2, #445231-C2; Mm-Vip-C2, # 415961-C2; Mm-Sst-C2, #404631-C2; Mm-Sst-C3, #404631-C3; Mm-Ndnf-C3, #447471-C3; Mm-Pvalb-C3, #421931-C3), followed by amplifying hybridization processes. DAPI was used as a nuclear stain. Prolong Gold Antifade (Thermo Scientific) was used to mount slides. Images were acquired on a confocal microscope (Zeiss LSM 880). EYFP-expressing cells, their distance from the pia and overlap with markers were quantified using a custom written MATLAB (MathWorks) script. Although the EGFP (anti-sense) probe labeled both mRNA and viral DNA, resulting in sparse puncta throughout the site of AAV2/5-EF1a-fDIO-EYFP-WPRE injection in ACx, cells that expressed mRNA (made possible via FLPo recombination) could be clearly distinguished from those that did not based on the intensity of expression. This was confirmed by the fact that an EGFP-sense probe, which only labels viral DNA, yielded only punctate labeling throughout the injection site but showed no strong labeling in individual cells. Anterograde transsynaptic tracing using AAV1 has been employed in a number of brain areas from both glutamatergic and GABAergic afferents to both excitatory and inhibitory targets (Beltramo and Scanziani, 2019; Benavidez et al., 2021; Lee et al., 2020; Trouche et al., 2019; Zingg et al., 2017; Zingg et al., 2020). The approach depends on high viral titers and signal amplification via DNA recombinases (Trouche et al., 2019; Zingg et al., 2017; Zingg et al., 2020). Although the precise molecular mechanisms are not fully understood, the available evidence indicates that AAV1 is trafficked down the axon, is not released from fibers of passage, and that transneuronal spread is strongly dependent on synaptic transmitter release (Zingg et al., 2017; Zingg et al., 2020). The efficacy of transsynaptic spread was estimated to be roughly equal for excitatory and inhibitory afferents and excitatory and inhibitory postsynaptic targets (Zingg et al., 2020). This method recapitulates established connectivity patterns (Zingg et al., 2020), and AAV1 labelled neurons are more likely to receive functional synaptic input from the afferents in question than their unlabeled neighbors (Trouche et al., 2019; Zingg et al., 2020). While AAV1 can also spread retrogradely along axons, this caveat does not apply to the data presented here since we identify overwhelmingly postsynaptic INs which do not extend their axons to long-range targets. These results indicate that the differences in neuronal labeling we observe using this technique are highly likely to derive from differences in synaptic connectivity.

### Histology

Mice were anesthetized i.p. with 300 mg/kg ketamine and 20 mg/kg xylazine (WDT) and transcardially perfused with 4% paraformaldehyde (PFA) in PBS. Brains were post-fixed overnight in 4% PFA at 4 °C and then stored in PBS. Coronal sections (60-100 μm thick) were cut using a Leica vibratome (VT1000S) and washed in PBS. Vibratome sections were permeabilized with 0.5% triton (Sigma) and then blocked in PBS-0.2% gelatin with 10% normal goat serum (Sigma), 0.2M glycine and 0.5% triton either overnight at 4 °C or at room temperature for 4 h. In cases where mouse primary antibodies would be used, 1:50 goat anti-mouse IgG antigen-binding fragments were included in the blocking solution (Jackson ImmunoResearch). Sections were incubated with primary antibodies in PBS-0.2% gelatin with 5% normal goat serum and 0.5% triton for 72 h at 4 °C. Primary antibodies used were the following: mouse anti-NeuN (1:500, RRID: AB_2298772, Merck Millipore) or rabbit anti-RFP (1:500, RRID: AB_591279, MBL). Sections were then washed in PBS with 0.5% triton and incubated, either overnight at 4 °C or at room temperature for 4 h, with fluorophore-conjugated secondary antibodies (1:1000, goat, Thermo Fisher Scientific) in PBS-0.2% gelatin with 5% normal goat serum and 0.5% triton. DAPI was used as a nuclear stain (5nM in PBS). Sections were mounted in Mowiol 4-88 (Polysciences) and imaged on a Zeiss confocal microscope (LSM 880). Brain regions were assigned using the Paxinos and Franklin’s mouse brain atlas (Paxinos and Franklin, 1997). For ZI axon quantifications in cortex, A1 and AuV were defined as regions in 500 μm blocks moving medially along the pia away from the rhinal fissure. L1 depth was calculated separately for both regions based on the DAPI signal. L1/L2 border was defined as the last bin before the DAPI fluorescence intensity (bin size 10 μm, as a function of depth from the pia) exceeded 1 standard deviation above the average of the first 80 μm for 2 consecutive bins. For retrograde labeling quantifications, ZI was identifiable from GAD2-nuclear-mCherry labeling. Labeled cells and overlap with mCherry signal were quantified manually from sections spanning the full anteroposterior extent of the ZI (from bregma, ~-1.22 until ~-2.92mm). For rabies tracing, mice were sacrificed 7 days post-RV injection. Perfused brains were cut in 60 μm coronal sections and counterstained with DAPI (30 min in 0.5 mg/ml, D1306, Thermo Fisher Scientific). To quantify the cell number per animal, every third section of the entire brain was scanned using Zen software (Zeiss) and cells were counted using a custom written MATLAB (MathWorks) script. To define the cell numbers in different brain regions, images were registered to the Allen Brain Atlas (Fürth et al., 2018). Only brain regions that revealed presynaptic cells in all mice are reported. The following histological images are compounds obtained by ‘stitching’ of different fields-of-view: Fig. 1, B and C; fig. 2, B and C.

### Acute brain slice recordings with optogenetics

35-40 days after virus injection, mice were anesthetized with isoflurane (5%) and decapitated into ice cold slicing solution containing (in mM): 93 NMDG, 2.5 KCl, 1.2 NaH_2_PO_4_, 30 NaHCO_3_, 20 HEPES, 25 glucose, 5 sodium ascorbate, 2 thiourea, 3 sodium pyruvate, 10 MgSO_4_ and 0.5 CaCl_2_ (titrated to pH 7.3-7.4 with HCl 1M, 310 mOsm). Coronal slices (350 μm thick) were prepared on a vibratome (Leica VT 1200S) and incubated in slicing solution at 33 °C for 15 minutes (Ting et al., 2014). Before the start of recordings, slices were incubated >45 minutes at room temperature in standard ACSF containing (in mM): 125 NaCl, 3 KCl, 1.25 NaH_2_PO_4_, 26 NaHCO_3_, 10 glucose, 1 MgCl_2_ and 2 CaCl_2_ (310 mOsm). Solutions were continuously bubbled with carbogen gas (95% O_2_, 5% CO_2_). Slice recordings were performed at 31-34 °C. Cells were visualized using differential interference contrast microscopy (Scientifica slice scope), a water immersion objective (40x, 0.8 N.A., Olympus LUMPLFLN) and a CCD camera (Hamamatsu C11440 ORCA-flash4.0). Fluorophore excitation was performed with LEDs through the objective (Cool LED). Whole-cell recordings were performed with a Multiclamp 700B amplifier (Axon Instruments), low-pass filtered at 3 kHz and digitized at 20 kHz (Digidata 1550 and pClamp software, Molecular Devices). Series resistance compensation was set at 80%. Patch pipettes (4-6 MΩ) were pulled from borosilicate capillaries with filaments and filled with intracellular solution (in mM): 140 K-gluconate, 10 KCl, 10 HEPES, 4 Na-phosphocreatine, 4 ATP-Mg, 0.4 GTP (pH 7.3, 290 mOsm). For ChR2-assisted circuit mapping (Petreanu et al., 2007), optogenetic stimulation of ZI axons was applied to superficial cortical layers using a 488 nm LED at 66.8 mW/mm^2^ irradiance. Inhibitory postsynaptic currents in response to a 5 ms light pulse were recorded in neurons clamped at 0 mV. In some experiments, SR 9553 hydrobromide (10 μM, Tocris) was applied to block GABAAR-mediated currents. L1 neurons and L2/3 pyramidal neurons were identified based on soma location and morphology.

### Discriminative threat conditioning paradigm (DTC) in freely-behaving animals with chemogenetics

Freely-behaving DTC experiments consisted of 3 sessions, each separated by 24hrs. In each session, mice were presented with 2 sounds (conditioned stimuli, CS), which consisted of trains of 500 ms frequency-modulated (FM) sweeps (logarithmically modulated between 5-15 kHz in the upwards direction, or between 20-10 kHz in the downwards direction, with 50 ms rise/fall) delivered at 1 Hz at 75 decibel (dB) sound pressure level (SPL) at the speaker (designed in RPvdsEx, processed by RZ6 and delivered by MF1 speakers, Tucker-Davis Technologies). All sessions took place in freely-behaving contexts. In session 1 (habituation) and in session 3 (memory recall), sounds were presented alone. In session 2 (acquisition), the CS+ co-terminated with a mild foot-shock (unconditioned stimulus, US), while the CS- did not. To induce local silencing of hM4D(Gi)-expressing (in control animals, EGFP/EYFP-expressing) axons prior to the conditioning session, animals were briefly anesthetized with isofluorane and 200nl of 3 nM clozapine-N-oxide (CNO, #C0832, Merck) was infused bilaterally via implanted cannulae in ACx using a Hamilton syringe (10 μl, 701N, Merck) and Nanoject stereotaxic syringe pump (Chemyx) (Meira et al., 2018; Morse et al., 2020). 15-30 min after recovery, animals were placed in the conditioning chamber and the session began. Up-FM-sweeps and down-FM-sweeps were used as CS+ and CS- in a counterbalanced fashion between animals. In habituation, CS+ and CS- FM-sweep trains lasting 30 s were presented in an interleaved fashion, 4 times each, starting with the CS+. In conditioning, CS+ and CS- FM-sweep trains lasting 10 s were presented in an interleaved fashion, 15 times each, starting with the CS+, with pseudorandom inter-trial intervals (ITIs) between 20 s and 180 s. The onset of the last FM-sweep of each CS+ coincided with the onset of a foot-shock US, delivered via the floor (1 s, 0.6 mA AC, Coulbourn Precision Animal Shocker, Coulbourn Instruments). Habituation took place in the conditioning context, while cued (auditory) memory recall was performed in a new context. The contexts were cleaned before and after each session with 70% ethanol and 0.2% acetic acid, respectively. In recall, the CS- was presented 4 times followed by 4 presentations of the CS+, both lasting 30 s each and presented at pseudorandom time intervals. This is a standard paradigm design for research on threat memory (Abs et al., 2018; Ciocchi et al., 2010; Courtin et al., 2014; Dalmay et al., 2019; Letzkus et al., 2011). As a measure of associative threat memory, freezing behavior was recorded using a webcam (HD C270, Logitech) and scored using a custom written MATLAB script (Abs et al., 2018; Castaño-Díez, 2018; Dalmay et al., 2019). Mice were considered to be freezing if no movement was detected for 2 s, and the measure was expressed as a percentage of time spent freezing. For cued memory recall, habituation and conditioning (Fig. 2G), freezing was quantified in 30 s intervals, starting at the onset of every CS. Baseline freezing was calculated during 30 s intervals randomly sampled in the first 3 minutes of each session, before any CS was presented. Discrimination index was calculated as CS+ freezing/(CS- freezing + CS+ freezing). For contextual memory recall, mice were placed in the conditioning context for 5 min, and freezing was quantified throughout the whole session.

### Discriminative threat conditioning (DTC) paradigm in head-fixation

Head-fixed DTC experiments consisted of 3 sessions, each separated by 24hrs. In each session, mice were presented with 2 sounds (CS), which consisted of trains of 500 ms FM sweeps (logarithmically modulated between 5-15 kHz in the upwards direction, or between 20-10 kHz in the downwards direction, with 50 ms rise/fall) delivered at 1 Hz at 75 dB SPL at the speaker (designed in RPvdsEx, processed by RZ6 and delivered by MF1 speakers, Tucker-Davis Technologies). All sessions took place in head-fixation under the microscope. In all sessions, FM-sweep trains lasting 10s were presented in an alternating fashion, 15 times each, with 70s inter-trial intervals (plus random interruptions required to adjust the microscope). In session 1 (habituation) and in session 3 (memory recall), sounds were presented alone. In session 2 (acquisition), the CS+ co-terminated with a mild tail-shock (unconditioned stimulus, US; 1s, 0.4mA, 10Hz; delivered with ISO-STIM 01D (NPI Electronic), using Pulse Pal v2 (Sanworks) as a pulse train generator), while the other (CS-) did not. Shocks were applied to the tail using adhesive electrode tabs (#TER-MXT-1334, TerniMed) which were cut in half length-wise, and then placed approximately 1 cm apart at the center of the tail (Dolensek et al., 2020). A lightweight, isolated cable with crocodile clamps connected the electrodes to the ISO-STIM 01D box. In sessions 1 and 3, the CS- was presented first, while in session 2, the CS+ was presented first. Up-FM-sweeps and down-FM-sweeps were used as CS+ and CS- in a counterbalanced fashion between animals. In all sessions, eye pupil diameter was recorded under infrared illumination (620 nm LED) at 16 Hz (Basler acA1920-25um camera) using custom-written software (Kretschmer, 2020a) to assess associative threat memory to the CS- and CS+. Pupil dilation in response to the CSs was calculated as the integral of ΔD/D0 in a 15 s time window for habituation and recall, where D0 is the mean pupil diameter within 2 s before sound-train onset and ΔD=D(t)-D0, where D(t) is the diameter at time t. In the acquisition session, we used a 9.5s time window since the tail-shock caused a rapid constriction of the pupil (Larsen and Waters, 2018; Silva et al., 2016). Pupil dilation in response to the shock was calculated the same way as for the CSs, but in a 7.5s time window starting 3.5s after the shock ended. To account for the different time windows between sessions, values were normalized to a time interval of 1 s. Discrimination index was calculated as CS+/(CS- + CS+) (Pollack et al., 2018). In habituation and acquisition, mice were head-fixed in a body tube made of plastic and lined with electrical insulating tape, and a plastic dish filled with 70% ethanol was placed inside the microscope chamber. In order to change the behavioral context during recall, a body tube made of metal was used, and a plastic dish filled with 0.2% acetic acid was placed inside the microscope chamber. Pseudoconditioning was conducted like DTC except that during acquisition, CSs and USs were presented in an explicitly unpaired fashion (15 presentations each for CS1, CS2 and shock). In this case, none of the CSs elicited any associative memory, and they were therefore pooled for analysis (Abs et al., 2018; Dalmay et al., 2019; Pardi et al., 2020).

### *In vivo* calcium imaging

Secondary auditory cortex was localized with intrinsic imaging under 1 % isoflurane anesthesia as in previous work (Abs et al., 2018; Dalmay et al., 2019; Pardi et al., 2020). Mice were water restricted and water delivery was used to facilitate habituation to handling and head-fixation on 6 consecutive days (4 days in the recording setup). Water was administered *ad libitum* before the experiments in head-fixation. Calcium imaging was performed with a resonant scanning microscope (Bruker Investigator). The femtosecond laser (Spectra Physics InSight) was tuned to 920 nm to excite axonGCaMP6s and tdTomato at an average excitation power under the objective (Nikon 16x, 0.8 N.A., 3 mm WD) of 20-25 mW. Images (512×512 pixels, 140×140 to 166×166 μm^2^) were acquired at 30 Hz in L1. Image acquisition, sound delivery and the camera for pupil tracking were controlled using custom written software (Kretschmer, 2020b; Kretschmer and Unzué, 2020). For image analysis, acquired time series were first corrected for motion, taking tdTomato images as a reference and using a custom MATLAB code (Kretschmer and Castaño-Díez, 2020) where data was temporally binned every 2 frames. Based on the average response in the red channel (tdTomato) to the CS+ in conditioning trials, or to the tail-shock alone in pseudoconditioning trials, a window of exclusion surrounding the tail shock delivery was defined to account for the fact that some boutons could not be motion corrected during this period (exclusion window from 1s before, until 3.5s after shock delivery). Regions of interest (ROIs) were selected in Fiji from the axonGCaMP6s fluorescence time average of the entire session. The following pipeline in Fiji allowed selection of both small and large boutons in a semi-automated way: first, the average axonGCaMP6s fluorescence image was filtered (maximum filter, 1 pixel radius); the resulting image was thresholded with the IsoData algorithm, keeping the ~5% highest intensity pixels; the resulting binary image was segmented with the *watershed* command; ROIs were obtained by applying the particle analysis, excluding particles on edges. Finally, ROIs were corrected manually in cases where more than 1 bouton was included inside an ROI, or in cases where ROIs delineated an axon segment without any bouton. Small boutons missed by the thresholding were also added manually. To superimpose frames from all sessions, axonGCaMP6s images were first translated and cropped in Fiji. ROIs were then obtained for each session independently. ROI sets from all three sessions were overlaid, and overlaying ROIs were assigned as *paired boutons*. Only paired boutons were used in the analysis. Average GCaMP6 fluorescence was measured for each ROI and frame, and data was subsequently analyzed in MATLAB. ROIs that displayed flat fluorescent traces without any calcium transients in the entire session were discarded. Traces were normalized either as ΔF/F_0_ or as z-score=ΔF/σ, where ΔF= F_(t)_-F_0_, with F_(t)_ being the fluorescence at a given time t, and F_0_ and σ the mean fluorescence before sound-train in each trial and its standard deviation, respectively. Boutons were considered sound responsive if they displayed significant activity that started no later than 10 s after sound-train onset (averaged z-score≥1.96 or ≤-1.645). Responses during FM-sweep sound-train were measured as the integral of ΔF/F_0_ from sound-train onset to the end of the sound train (10 s sound-trains), except for conditioning, where only the first 9 s were measured to exclude shock-related motion artefacts or responses. To account for the different time windows between sessions, values were normalized to a time interval of 1 s. Responses following shock delivery in conditioning or pseudoconditioning sessions were measured as the integral of ΔF/F_0_ starting from 3.5s after shock delivery, and then for 5s. DTC boutons were segregated into two paired populations across sessions based on whether their mean CS response (ΔF/F_0_*s) in recall was above (‘positive response in recall’, PR boutons) or below (‘negative response in recall’, NR boutons) zero. Latency to response peak was calculated as the time from CS onset to the global response peak within 9s for all 3 DTC sessions, in consideration of the shock during the conditioning session. For correlation with memory strength, we compared pupil and bouton response in habituation or recall for each mouse and stimulus (CS+, CS-, CS1/2). This was done independently for PR and NR boutons, and additionally the absolute value of these changes were added together to obtain the ‘absolute bouton response’.

### Stimulus information in individual boutons

All computational analyses were performed in Python, using the NumPy (Harris et al., 2020), SciPy (Virtanen et al., 2020) and Scikit-learn (Pedregosa et al., 2012) libraries. The amount of stimulus-specific information in the response of individual boutons was quantified using the sensitivity index, defined as:

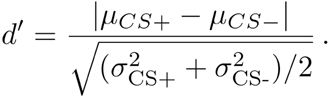

Here *μ_CS+_* and *μ_CS-_* are a bouton’s mean response during the CS+ and CS- presentation, averaged over the duration of the stimulus and all trials in which stimulus *s* was presented:

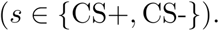

The variance of the responses across trials, after averaging across the stimulus window for each trial, is:

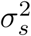

### Decoding from populations of boutons

To assess how much stimulus information is present on the population level, we trained linear decoders to predict stimulus identity from synaptic activity. For each trial, we collected the time-varying population response during stimulus presentation, binned in time windows of 1s. We excluded the last (shock) bin. Each stimulus was presented 15 times for 10s, yielding 9 time bins for each of the 2×15 trials. We decoded the stimuli using L2-regularized logistic regression, using leave-two-trial out cross-validation to avoid overfitting and balance the number of training samples per stimulus. The remaining 28×9 training samples were randomly divided into 10 cross-validation folds, which were used to determine the regularization strength of logistic regression. The candidate regularization strengths were 10 values, equally spaced on a logarithmic scale between 1e-4 and 1e4 (the Scikit-learn default). These values determine the inverse regularization strength, such that smaller values correspond to stronger regularization. For optimal regularizing strength, we tested the classifier on the 2 left-out trials, repeating this cross-validation procedure by leaving out all pairs of consecutive trials:

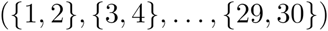

The average accuracy (% of time frames correct) over all of these test trials is reported. To estimate chance level performance, we repeatedly trained classifiers after randomly permuting the stimulus labels of individual time frames. Decoders were trained on data from individual mice and on pooled data, for which we concatenated the responses from all mice for corresponding trials in the protocol. For the pooled data, we first used principal component analysis (PCA) to reduce the data from the total number of boutons down to 200 dimensions, preserving approximately 90% of the variance. This sped up the analyses without qualitatively changing the results. We did not reduce the dimensionality before decoding from individual animals. To test if the higher decoding accuracy after conditioning compared to pseudoconditioning was due to a larger number of recorded boutons, we subsampled boutons from the conditioned mice. Specifically, decoders were trained on 100 random subsets of 535 boutons from conditioned mice, since this was the total number of boutons from all mice in our pseudoconditioning dataset. To assess whether differences in decoding accuracy were significant, bootstrapping was used to estimate the sampling distribution of the decoding accuracy. Specifically, we trained decoders on 1000 resamples (with replacement) of all boutons.

### Dynamics of population vectors over learning

To assess how stimulus-evoked population activity changes over the course of learning, we computed angles between population vectors at different moments in time and for different stimulus conditions. The change from habituation to recall was quantified as the ‘learning angle’ between the mean population vectors for each session. The mean vectors were computed as the average population response, averaged across all trials and stimulus bins (again excluding the shock window), for each stimulus separately. The dynamics of the population vector during acquisition was quantified by the angle between the average population vector during habituation (or recall) and the single-trial population vectors during acquisition. The single-trial vectors were computed as the time-averaged response during stimulus presentation. To assess whether the differences in the population vector from one day to the next (i.e., between habituation and the beginning of acquisition, and the end of acquisition and recall) are due to trial-to-trial fluctuations or due to representational changes, we quantified the trial-to-trial variability during habituation and recall. To this end, we computed the average of the angle between the mean population vector during habituation (or recall) and the single-trial habituation (recall) vectors. To avoid underestimating trial-to-trial variability, we computed the angle between the population vector of trial t and the average vector from all trials other than trial *t*.

### Response variability and correlations across boutons

We observed that conditioning generated both positive and negative stimulus responses, thereby increasing the response variability across boutons. To quantify this effect, we computed the standard deviation across boutons of their trial-averaged responses, for each moment during stimulus presentation and for each mouse. We also analyzed the structure of trial variability, by computing noise correlations between all pairs of simultaneously imaged boutons. To this end, we first computed the time-averaged response for each bouton and trial, before computing Pearson correlation coefficients. Correlation coefficients were then pooled across mice. We left out the first two trials, since the initial stimulus response on those trials led to unusually high noise correlations (of almost 1) for many boutons.

### Statistical analysis

was performed using GraphPad Prism and MATLAB. Data were considered normally distributed if Shapiro-Wilk, D’Agostino & Pearson and KS tests were passed. According to this result, and depending on whether data was paired or not, comparisons were performed using the following parametric or non-parametric tests: For two-group comparisons, 2-tailed t-test (normal distribution, non-paired), 2-tailed paired t-test (normal distribution, paired), 2-tailed Mann-Whitney test (non-normal distribution, non-paired) and 2-tailed Wilcoxon test (non-normal distribution, paired). For three-group comparisons, One-way ANOVA (normal distribution, non-paired) or One-way repeated measures (RM) ANOVA (normal distribution, paired) followed by Tukey’s multiple comparisons test, and Kruskal-Wallis test (non-normal distribution, non-paired) or Friedman test (non-normal distribution, paired) followed by Dunn’s multiple comparisons test. Two-way RM ANOVA followed by Sidak’s and Tukey’s multiple comparisons tests were used to compare groups across more than one factor. Correlations were computed as Pearson coefficients. Statistical tests used in each instance are indicated in the figure legends. Results are reported as: n.s. (not significant) p>0.05, * p<0.05, ** p<0.01, *** p<0.001, **** p<0.0001.

### Code availability

Codes necessary to understand and assess the conclusions of the manuscript were archived in: Mouse freezing behavior analysis: https://doi.org/10.17617/1.8Q (Castaño-Díez, 2018)

Eye tracker plugin for PylonRecorder: http://doi.org/10.17617/1.8M (Kretschmer, 2020a)

PylonRecorder camera recording software for eye imaging: http://doi.org/10.17617/1.8N (Kretschmer, 2020b)

AudioGameGUI for sounds, camera and microscope synchronization: http://doi.org/10.17617/1.8O (Kretschmer and Unzué, 2020)

Motion correction for calcium imaging data: https://doi.org/10.17617/1.8P (Kretschmer and Castaño-Díez, 2020)

Computational analysis: https://github.com/sprekelerlab/long-range-inhibition (Keijser, 2022)

**Table S1.**
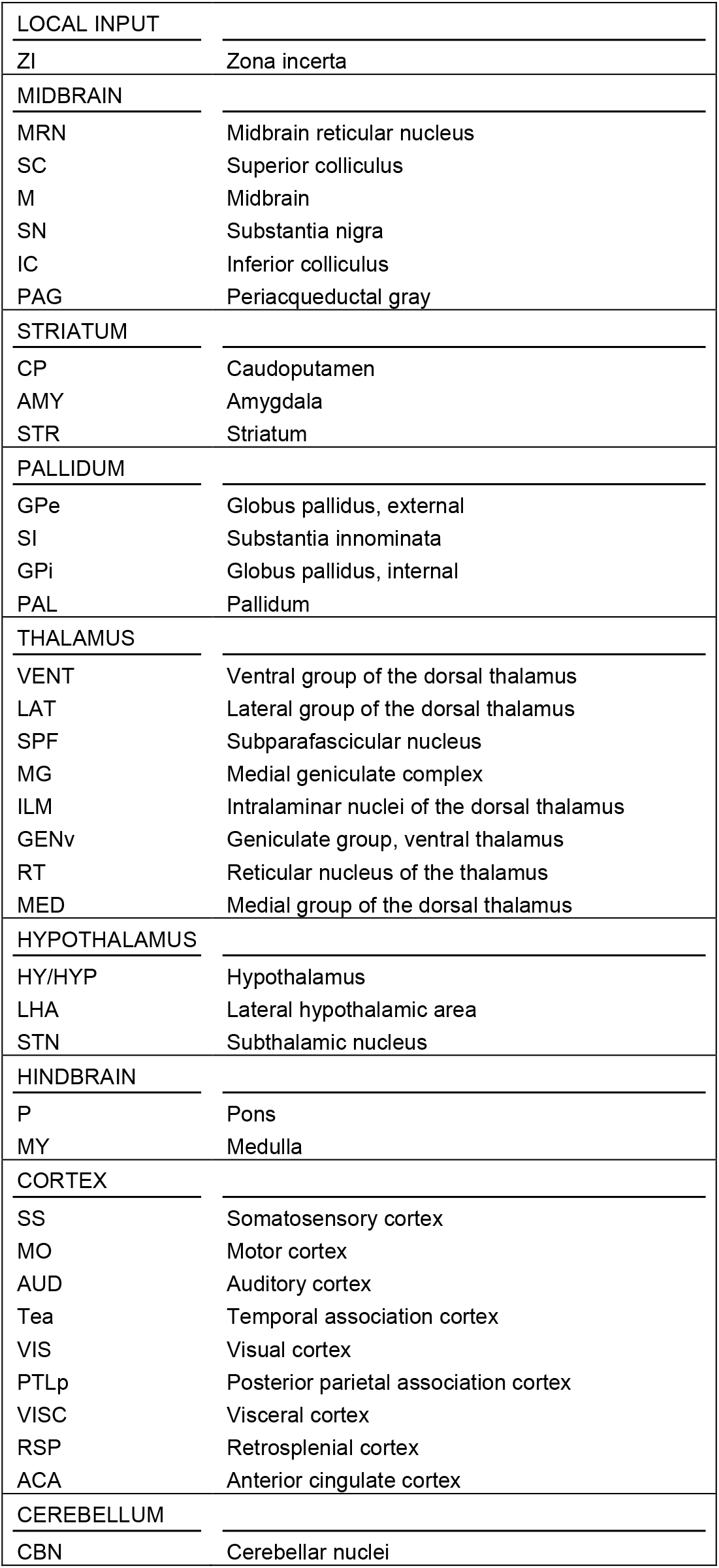
Abbreviations for brain structures.

## Notes

### Competing Interest Statement

The authors have declared no competing interest.

## References and Notes

Abs, E., Poorthuis, R.B., Apelblat, D., Muhammad, K., Pardi, M.B., Enke, L., Kushinsky, D., Pu, D.L., Eizinger, M.F., Conzelmann, K.K., et al. (2018). Learning-Related Plasticity in Dendrite-Targeting Layer 1 Interneurons. Neuron 100, 684–699 e686.

Ahmadlou, M., Houba, J.H.W., van Vierbergen, J.F.M., Giannouli, M., Gimenez, G.A., van Weeghel, C., Darbanfouladi, M., Shirazi, M.Y., Dziubek, J., Kacem, M., et al. (2021). A cell type-specific cortico-subcortical brain circuit for investigatory and novelty-seeking behavior. Science 372.

Basu, J., Zaremba, J.D., Cheung, S.K., Hitti, F.L., Zemelman, B.V., Losonczy, A., and Siegelbaum, S.A. (2016). Gating of hippocampal activity, plasticity, and memory by entorhinal cortex long-range inhibition. Science 351, aaa5694.

Beltramo, R., and Scanziani, M. (2019). A collicular visual cortex: Neocortical space for an ancient midbrain visual structure. Science 363, 64–69.

Benavidez, N.L., Bienkowski, M.S., Zhu, M., Garcia, L.H., Fayzullina, M., Gao, L., Bowman, I., Gou, L., Khanjani, N., Cotter, K.R., et al. (2021). Organization of the inputs and outputs of the mouse superior colliculus. Nature communications 12, 4004.

Broussard, G.J., Liang, Y., Fridman, M., Unger, E.K., Meng, G., Xiao, X., Ji, N., Petreanu, L., and Tian, L. (2018). In vivo measurement of afferent activity with axon-specific calcium imaging. Nat Neurosci 21, 1272–1280.

Castaño-Díez, D. (2018). Freezing Analysis, GitHub. doi:10.17617/17611.17618Q.

Ceballo, S., Piwkowska, Z., Bourg, J., Daret, A., and Bathellier, B. (2019). Targeted Cortical Manipulation of Auditory Perception. Neuron 104, 1168–1179.e1165.

Chen, J., and Kriegstein, A.R. (2015). A GABAergic projection from the zona incerta to cortex promotes cortical neuron development. Science 350, 554–558.

Ciocchi, S., Herry, C., Grenier, F., Wolff, S.B., Letzkus, J.J., Vlachos, I., Ehrlich, I., Sprengel, R., Deisseroth, K., Stadler, M.B., et al. (2010). Encoding of conditioned fear in central amygdala inhibitory circuits. Nature 468, 277–282.

Courtin, J., Chaudun, F., Rozeske, R.R., Karalis, N., Gonzalez-Campo, C., Wurtz, H., Abdi, A., Baufreton, J., Bienvenu, T.C., and Herry, C. (2014). Prefrontal parvalbumin interneurons shape neuronal activity to drive fear expression. Nature 505, 92–96.

Dalmay, T., Abs, E., Poorthuis, R.B., Hartung, J., Pu, D.L., Onasch, S., Lozano, Y.R., Signoret-Genest, J., Tovote, P., Gjorgjieva, J., et al. (2019). A Critical Role for Neocortical Processing of Threat Memory. Neuron 104, 1180–1194 e1187.

Dejean, C., Courtin, J., Karalis, N., Chaudun, F., Wurtz, H., Bienvenu, T.C., and Herry, C. (2016). Prefrontal neuronal assemblies temporally control fear behaviour. Nature 535, 420–424.

Dolensek, N., Gehrlach, D.A., Klein, A.S., and Gogolla, N. (2020). Facial expressions of emotion states and their neuronal correlates in mice. Science 368, 89–94.

Doron, G., Shin, J.N., Takahashi, N., Druke, M., Bocklisch, C., Skenderi, S., de Mont, L., Toumazou, M., Ledderose, J., Brecht, M., et al. (2020). Perirhinal input to neocortical layer 1 controls learning. Science 370.

Fürth, D., Vaissière, T., Tzortzi, O., Xuan, Y., Märtin, A., Lazaridis, I., Spigolon, G., Fisone, G., Tomer, R., Deisseroth, K., et al. (2018). An interactive framework for whole-brain maps at cellular resolution. Nat Neurosci 21, 139–149.

Gambino, F., Pages, S., Kehayas, V., Baptista, D., Tatti, R., Carleton, A., and Holtmaat, A. (2014). Sensory-evoked LTP driven by dendritic plateau potentials in vivo. Nature 515, 116–119.

Harris, C.R., Millman, K.J., van der Walt, S.J., Gommers, R., Virtanen, P., Cournapeau, D., Wieser, E., Taylor, J., Berg, S., Smith, N.J., et al. (2020). Array programming with NumPy. Nature 585, 357–362.

Herry, C., and Johansen, J.P. (2014). Encoding of fear learning and memory in distributed neuronal circuits. Nat Neurosci 17, 1644–1654.

Kaifosh, P., Lovett-Barron, M., Turi, G.F., Reardon, T.R., and Losonczy, A. (2013). Septo-hippocampal GABAergic signaling across multiple modalities in awake mice. Nat Neurosci 16, 1182–1184.

Keijser, J. (2022). Population-level analysis of long-range inhibition in neocortical layer 1, GitHub. https://github.com/sprekelerlab/long-range-inhibition.

Kretschmer, F. (2020a). EyeTracker, MPCDF Gitlab. doi:10.17617/17611.17618M.

Kretschmer, F. (2020b). PylonRecorder, MPCDF Gitlab. doi:10.17617/17611.17618N.

Kretschmer, F., and Castaño-Díez, D. (2020). MotionCorrection, MPCDF Gitlab. doi:10.17617/17611.17618P.

Kretschmer, F., and Unzué, D. (2020). AudioGameGUI, MPCDF Gitlab. doi:10.17617/17611.17618O.

Larsen, R.S., and Waters, J. (2018). Neuromodulatory Correlates of Pupil Dilation. Frontiers in neural circuits 12, 21.

Lee, J., Wang, W., and Sabatini, B.L. (2020). Anatomically segregated basal ganglia pathways allow parallel behavioral modulation. Nat Neurosci 23, 1388–1398.

Letzkus, J.J., Wolff, S.B., and Luthi, A. (2015). Disinhibition, a Circuit Mechanism for Associative Learning and Memory. Neuron 88, 264–276.

Letzkus, J.J., Wolff, S.B., Meyer, E.M., Tovote, P., Courtin, J., Herry, C., and Luthi, A. (2011). A disinhibitory microcircuit for associative fear learning in the auditory cortex. Nature 480, 331–335.

Li, Z., Rizzi, G., and Tan, K.R. (2021). Zona incerta subpopulations differentially encode and modulate anxiety. Sci Adv 7, eabf6709.

Liu, K., Kim, J., Kim, D.W., Zhang, Y.S., Bao, H., Denaxa, M., Lim, S.A., Kim, E., Liu, C., Wickersham, I.R., et al. (2017). Lhx6-positive GABA-releasing neurons of the zona incerta promote sleep. Nature 548, 582–587.

Makino, H., and Komiyama, T. (2015). Learning enhances the relative impact of top-down processing in the visual cortex. Nat Neurosci 18, 1116–1122.

Masri, R., Quiton, R.L., Lucas, J.M., Murray, P.D., Thompson, S.M., and Keller, A. (2009). Zona incerta: a role in central pain. J Neurophysiol 102, 181–191.

Meira, T., Leroy, F., Buss, E.W., Oliva, A., Park, J., and Siegelbaum, S.A. (2018). A hippocampal circuit linking dorsal CA2 to ventral CA1 critical for social memory dynamics. Nature communications 9, 4163.

Melzer, S., and Monyer, H. (2020). Diversity and function of corticopetal and corticofugal GABAergic projection neurons. Nat Rev Neurosci 21, 499–515.

Morse, A.K., Leung, B.K., Heath, E., Bertran-Gonzalez, J., Pepin, E., Chieng, B.C., Balleine, B.W., and Laurent, V. (2020). Basolateral Amygdala Drives a GPCR-Mediated Striatal Memory Necessary for Predictive Learning to Influence Choice. Neuron 106, 855–869.e858.

Ossowska, K. (2020). Zona incerta as a therapeutic target in Parkinson’s disease. J Neurol 267, 591–606.

Pardi, M.B., Vogenstahl, J., Dalmay, T., Spano, T., Pu, D.L., Naumann, L.B., Kretschmer, F., Sprekeler, H., and Letzkus, J.J. (2020). A thalamocortical top-down circuit for associative memory. Science 370, 844–848.

Paxinos, G., and Franklin, K.B. (1997). The mouse brain in stereotaxic coordinates. 2001. Brook, New York, USA), respectively Each experiment lasted around two and a half hours In the end of the experiment, a 5.

Pedregosa, F., Varoquaux, G., Gramfort, A., Michel, V., Thirion, B., Grisel, O., Blondel, M., Prettenhofer, P., Weiss, R., Dubourg, V., et al. (2012). Scikit-learn: Machine Learning in Python. Journal of Machine Learning Research 12.

Peron, S.P., Freeman, J., Iyer, V., Guo, C., and Svoboda, K. (2015). A Cellular Resolution Map of Barrel Cortex Activity during Tactile Behavior. Neuron 86, 783–799.

Petreanu, L., Huber, D., Sobczyk, A., and Svoboda, K. (2007). Channelrhodopsin-2-assisted circuit mapping of long-range callosal projections. Nat Neurosci 10, 663–668.

Petreanu, L., Mao, T., Sternson, S.M., and Svoboda, K. (2009). The subcellular organization of neocortical excitatory connections. Nature 457, 1142–1145.

Pollack, G.A., Bezek, J.L., Lee, S.H., Scarlata, M.J., Weingast, L.T., and Bergstrom, H.C. (2018). Cued fear memory generalization increases over time. Learning & memory (Cold Spring Harbor, NY) 25, 298–308.

Roelfsema, P.R., and Holtmaat, A. (2018). Control of synaptic plasticity in deep cortical networks. Nat Rev Neurosci 19, 166–180.

Saunders, A., Oldenburg, I.A., Berezovskii, V.K., Johnson, C.A., Kingery, N.D., Elliott, H.L., Xie, T., Gerfen, C.R., and Sabatini, B.L. (2015). A direct GABAergic output from the basal ganglia to frontal cortex. Nature 521, 85–89.

Shin, J.N., Doron, G., and Larkum, M.E. (2021). Memories off the top of your head. Science 374, 538–539.

Silva, B.A., Gross, C.T., and Gräff, J. (2016). The neural circuits of innate fear: detection, integration, action, and memorization. Learning & memory (Cold Spring Harbor, NY) 23, 544–555.

Taniguchi, H., He, M., Wu, P., Kim, S., Paik, R., Sugino, K., Kvitsiani, D., Fu, Y., Lu, J., Lin, Y., et al. (2011). A resource of Cre driver lines for genetic targeting of GABAergic neurons in cerebral cortex. Neuron 71, 995–1013.

Ting, J.T., Daigle, T.L., Chen, Q., and Feng, G. (2014). Acute brain slice methods for adult and aging animals: application of targeted patch clamp analysis and optogenetics. Methods in molecular biology (Clifton, NJ) 1183, 221–242.

Trageser, J.C., and Keller, A. (2004). Reducing the uncertainty: gating of peripheral inputs by zona incerta. J Neurosci 24, 8911–8915.

Trouche, S., Koren, V., Doig, N.M., Ellender, T.J., El-Gaby, M., Lopes-Dos-Santos, V., Reeve, H.M., Perestenko, P.V., Garas, F.N., Magill, P.J., et al. (2019). A Hippocampus-Accumbens Tripartite Neuronal Motif Guides Appetitive Memory in Space. Cell 176, 1393–1406 e1316.

Urbain, N., and Deschenes, M. (2007). Motor cortex gates vibrissal responses in a thalamocortical projection pathway. Neuron 56, 714–725.

Virtanen, P., Gommers, R., Oliphant, T.E., Haberland, M., Reddy, T., Cournapeau, D., Burovski, E., Peterson, P., Weckesser, W., Bright, J., et al. (2020). SciPy 1.0: fundamental algorithms for scientific computing in Python. Nature methods 17, 261–272.

Zhang, X., and van den Pol, A.N. (2017). Rapid binge-like eating and body weight gain driven by zona incerta GABA neuron activation. Science 356, 853–859.

Zhou, M., Liu, Z., Melin, M.D., Ng, Y.H., Xu, W., and Sudhof, T.C. (2018). A central amygdala to zona incerta projection is required for acquisition and remote recall of conditioned fear memory. Nat Neurosci 21, 1515–1519.

Zingg, B., Chou, X.L., Zhang, Z.G., Mesik, L., Liang, F., Tao, H.W., and Zhang, L.I. (2017). AAV-Mediated Anterograde Transsynaptic Tagging: Mapping Corticocollicular Input-Defined Neural Pathways for Defense Behaviors. Neuron 93, 33–47.

Zingg, B., Peng, B., Huang, J., Tao, H.W., and Zhang, L.I. (2020). Synaptic Specificity and Application of Anterograde Transsynaptic AAV for Probing Neural Circuitry. J Neurosci 40, 3250–3267.

